# Automated identification of small molecules in cryo-electron microscopy data with density- and energy-guided evaluation

**DOI:** 10.1101/2024.11.20.623795

**Authors:** Andrew Muenks, Daniel P. Farrell, Guangfeng Zhou, Frank DiMaio

## Abstract

Methodological improvements in cryo-electron microscopy (cryoEM) have made it a useful tool in ligand-bound structure determination for biology and drug design. However, determining the conformation and identity of bound ligands is still challenging at the resolutions typical for cryoEM. Automated methods can aid in ligand conformational modeling, but current ligand identification tools — developed for X-ray crystallography data — perform poorly at resolutions common for cryoEM. Here, we present EMERALD-ID, a method capable of docking and evaluating small molecule conformations for ligand identification. EMERALD-ID identifies 43% of common ligands exactly and identifies closely related ligands in 66% of cases. We then use this tool to discover possible ligand identification errors, as well as previously unidentified ligands. Furthermore, we show EMERALD-ID is capable of identifying ligands from custom ligand libraries of various small molecule types, including human metabolites and drug fragments. Our method provides a valuable addition to cryoEM modeling tools to improve small molecule model accuracy and quality.

## INTRODUCTION

Over the past decade, cryo-electron microscopy (cryoEM) has become widely used in macromolecular structure determination as advancements in both data collection and data processing have improved map resolutions. As EM data approaches atomic^1,2^ and near-atomic resolutions, protein-small molecule interactions are observable, leading to an increase in ligands modeled in cryoEM structures^3^) and the use of cryoEM in drug discovery^4^. Despite these advancements in data resolution, the lower resolution of typical cryoEM maps means building models into cryoEM data is still difficult and error-prone for both proteins^5^ and ligands^6^.

Building on recent advances in protein structure prediction from machine learning, numerous tools exist for robustly building protein models into cryoEM data^7–10^. However, tools for modeling ligands are less well-developed. While ligand fitting tools are available^11–13^, no capable methods exist to accurately identify ligands at moderate resolution data. Current automated ligand identification methods — primarily developed for crystallographic data — rely on density map correlations^14^ or shape features of the maps^15,16^, leading to limited accuracy at resolutions worse than 3 Å. While deep-learning methods for protein structure prediction now promise sequence to structure prediction of ligands bound to structures^17,18^, they only determine the ligand conformation, not identity, and are unaware of EM map information.

In order to produce accurate small molecule models and determine ligand identity, additional information to map features must be used. Here, we present EMERALD-ID, a ligand identification tool for cryoEM data. EMERALD-ID utilizes the RosettaGenFF small molecule force field^19^, the EMERALD ligand fitting method^12^, and a linear regression model combining estimated binding affinity and density correlation to discern ligand identities from a library. The accuracy of EMERALD-ID was evaluated on ligand-bound structures of common ligands in the Electron Microscopy Data Bank (EMDB)^20^, upon which we found 60 EMDB entries with a high-confidence EMERALD-ID solution different from the deposited model. Additionally, we searched deposited maps in the EMDB and identified 65 maps with plausible ligand omissions. Lastly, we show the robustness of EMERALD-ID by screening against large, diverse libraries of human metabolites and drug fragments.

## RESULTS

### Explanation of EMERALD-ID

An overview of EMERALD-ID is shown in Figure 1. EMERALD-ID takes a user-provided library of ligand identities, an EM density map, and a starting receptor model and docks all identities from the library into the EM map using EMERALD (Fig. 1A). To compare identities fairly, we created a linear regression model that considers ligand size, local resolution of the map around the binding pocket, and the density correlation of the receptor to predict an expected ligand density correlation and estimated binding affinity (ΔG) for a given map and ligand (Fig. 1B). The density correlations and ΔG values of all docked identities are compared to the expected values from the model to calculate a unitless Z-score. Once calculated, EMERALD-ID predicts the probability of each identity by a modified cross-entropy function and ranks the molecules (Fig. 1C).

**Figure 1.**
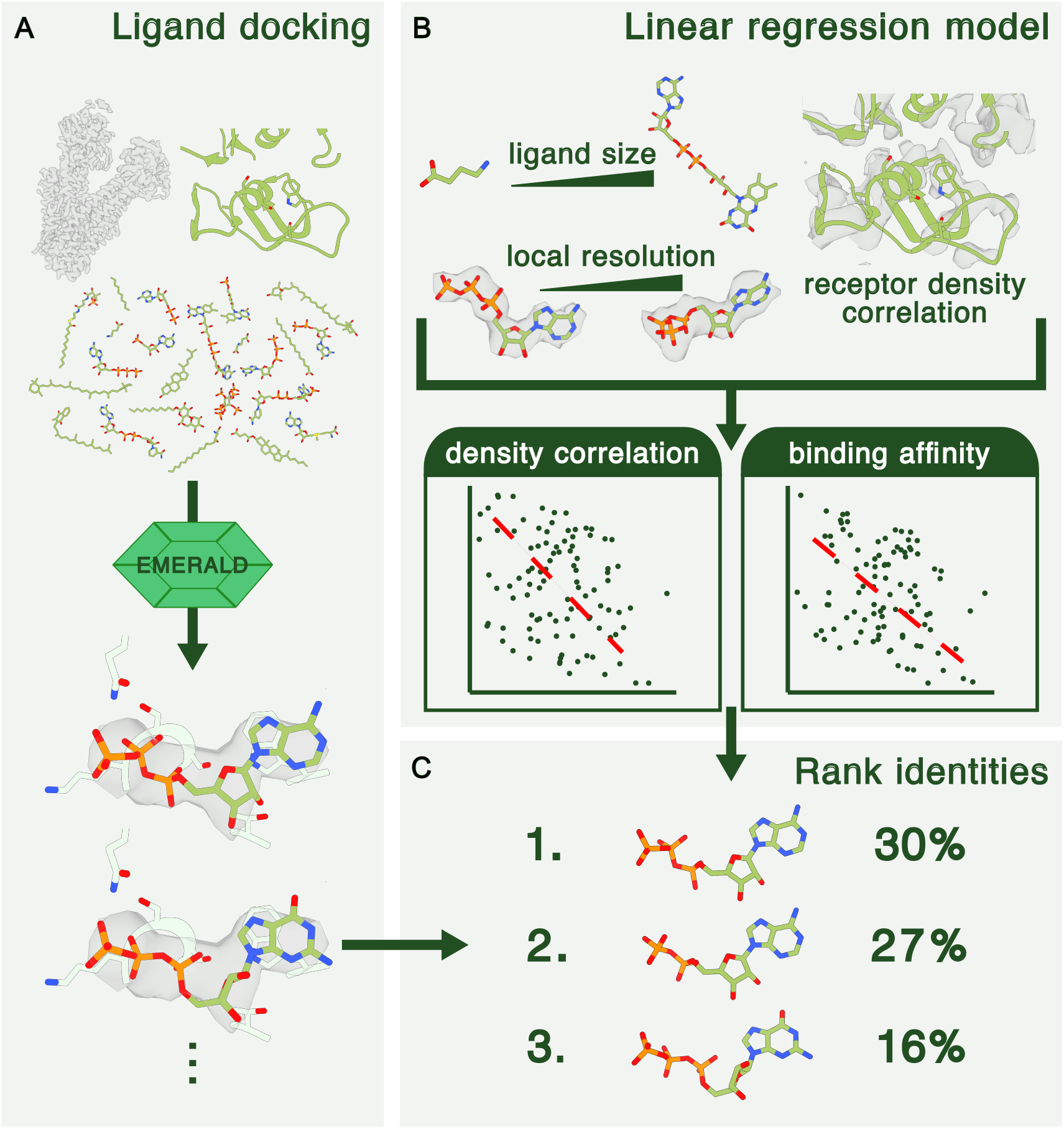
Overview of EMERALD-ID. (A) Identities from a provided library are fitted into the EM map with EMERALD. (B) A linear regression model takes features of ligand size, local resolution, and receptor fit into density and predicts expected density correlation and binding affinity values for a given identity-map pair. (C) Predicted values from the model are compared to calculated values from docked models to rank and assign probabilities to identities.

To test EMERALD-ID, we wanted to focus on scenarios modelers may experience during structure determination. First, we created a ligand library of thirty common ligands in cryoEM structures. With this library, we determined the accuracy of EMERALD-ID on deposited cryoEM structures, and furthermore, searched maps in the EMDB for unassigned density likely belonging to common ligands. Finally, we examined EMERALD-ID’s capabilities when considering a large endogenous ligand library, as well as its potential for fragment-based drug discovery.

### Evaluation of ligand identification in deposited structures

A common task in ligand identification is screening against a small library of common ligands. We decided to benchmark EMERALD-ID with this task. While the popular modeling suite Phenix provides a list of the most common ligands bound to macromolecular models, several of these ligands do not appear in any cryoEM solved structures. We set out to create our own list of common ligands solved with cryoEM. We settled on 30 common ligand identities to use for evaluation that encompass 38% of small molecule structures in cryoEM. This library included nucleotide substrates and cofactors like ATP and NADH as well as lipids like cholesterol and palmitate.

We gathered 1387 appearances of a common ligand identity from 1221 EMDB entries. All 30 ligands in the library were docked in the pocket of the first instance of the common ligand in the deposited structure. EMERALD-ID correctly ranked the deposited identity first for 43% of instances (Fig. 2A). Identification results were compared to phenix.ligand identification which determined the correct identity in 10% of cases (Fig. 2A). Our ability to correctly identify the ligand relied heavily on successfully docking the molecules. EMERALD-ID docked the native identity within 1 Å RMSD of the deposited structure for 39% of cases; in these cases it correctly identified the native ligand 68% of the time (Fig. 2B). Identification accuracy was also dependent on the local resolution of the binding pocket (Fig. 2C). We achieve an accuracy of 46% for all maps with 4.5 Å resolution or better, but accuracy plummeted at worse resolutions. This was unsurprising given the lack of detail in maps at low resolutions, and we previously showed that ligand fitting accuracy in EMERALD decreased at this same resolution^12^.

**Figure 2.**
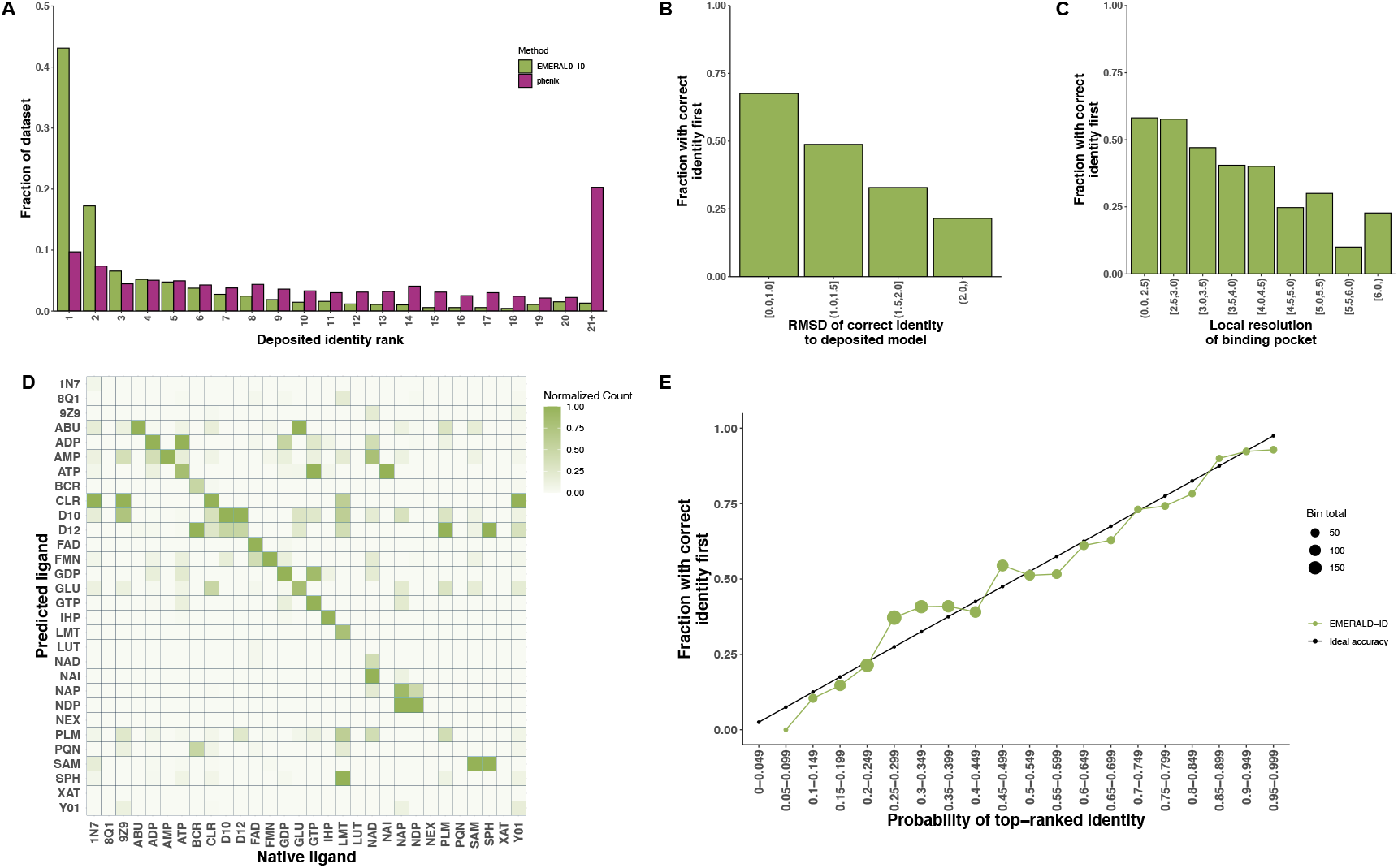
EMERALD-ID results from screening common ligand identities. (A) Fraction of the data for rank placements of the deposited identity for EMERALD-ID (green, n = 1387) and Phenix (purple, n = 1030). (B, C) Accuracy of ranking the deposited identity first by docking success (B) and local resolution (C). (D) Confusion matrix of common ligand identities. Ligands are labeled by their name in the Chemical Component Dictionary. Counts for each identity are normalized by column. (E) Comparison of predicted accuracy to true accuracy for EMERALD-ID.

In instances where EMERALD-ID did not identify the correct ligand, it often chose a closely related identity. In 66% of entries, the top ligand had a Tanimoto similarity coefficient greater than 0.75 to the deposited identity. EMERALD-ID often confused nucleotides that differed by phosphate length or base, which are ambiguous at medium to low map resolutions (Fig. 2D). For steroids and lipids, EMERALD-ID tends to favor smaller ligands within the class, e.g. cholesterol (CLR) vs. cholesterol hemisuccinate (Y01), which is expected given that the larger ligands likely have disordered regions that are not represented in the EM map.

Along with the rankings, we looked at the predicted probabilities provided by EMERALD-ID. The true accuracy of the common ligand screen closely matched the predicted accuracy of the top-ranked identity (Fig. 2E). Additionally, the predicted probabilities found possible identity corrections by highlighting high-confidence cases that do not rank the deposited identity first. Indeed, we found 60 “incorrect” cases that have a probability over 0.60. A common possible correction occurred between ATP and ADP. For example, in an ATP synthase^21^, the deposited structure placed an ATP molecule in the density, despite all 3 phosphates struggling to fit (Fig. 3A), while EMERALD-ID preferred an ADP molecule by both binding affinity and density fit (Fig. 3B). While the site is likely partially occupied by both identities, our metrics suggested that ADP was the more probable ligand.

**Figure 3.**
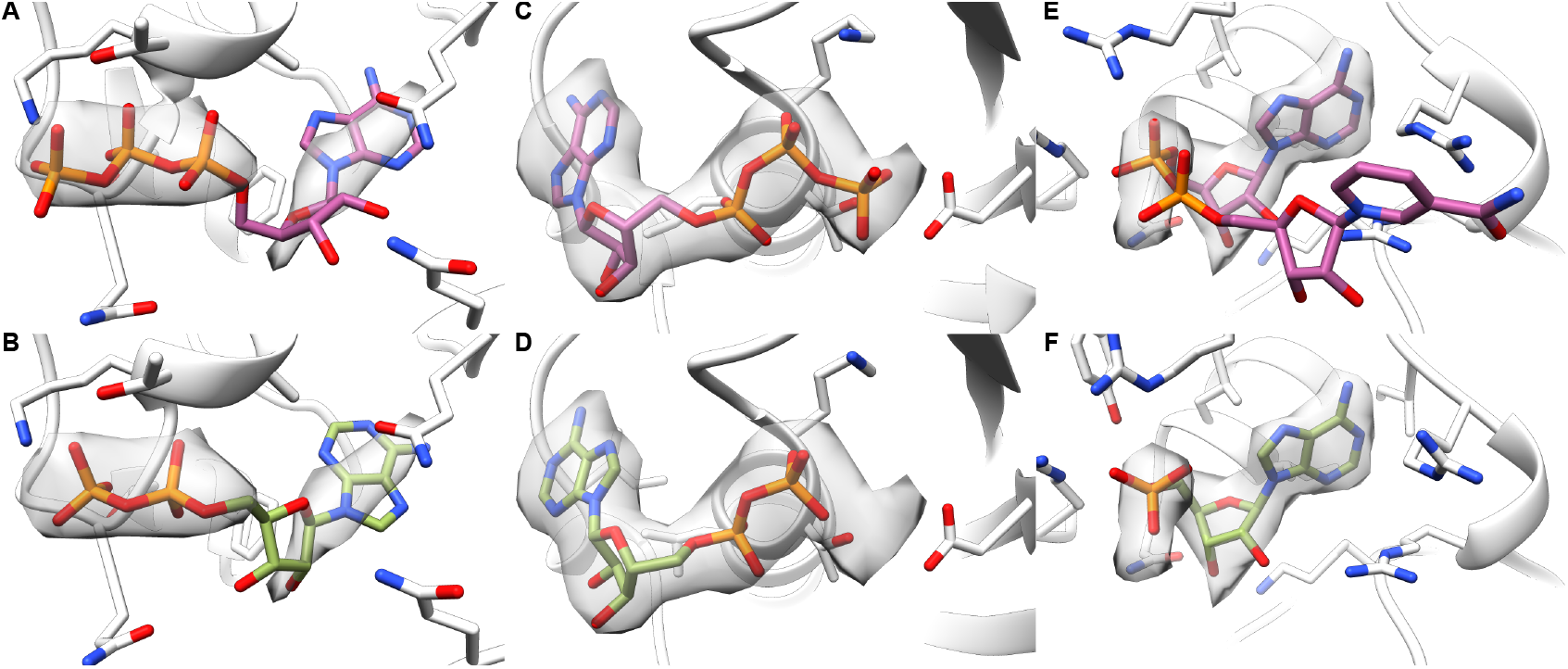
Examples of high-confidence EMERALD-ID identities different than the deposited model. (A, B) Deposited ATP molecule (A) is disfavored for an ADP molecule in EMERALD-ID (B) in an ATP synthase (EMDB: 21264, PDB: 6VOH). (C, D) Deposited ATP molecule (C) is replaced with an ADP molecule in EMERALD-ID (D) in the Ufd1/Npl4/Cdc48 complex (EMDB: 27273, PDB: 8DAR). (E, F) Deposited NAD^+^ molecule (E) in malic enzyme 2 (EMDB: 33145, PDB: 7XDE) is outscored by an AMP molecule (F).

Another example of mistaken nucleotide identity was found in a structure of the Ufd1/Npl4/Cdc4 complex^22^. In the deposited structure, the modeled ATP molecule satisfied the EM map, but in doing so placed the gamma phosphate near an aspartate residue (Fig. 3C). EMERALD-ID elected to avoid this repulsive clash and left a portion of the map unexplained with the top-ranked ADP molecule (Fig. 3D). Likely, the unexplained density belongs to a magnesium ion. Even if EMERALD-ID did not explicitly model the ion, it avoided overfitting into the density because the conformation does not fit energetically.

A final example includes a molecule that was too large for the observed density. In a malic enzyme 2 structure^23^, the nicotinamide moiety of the NAD^+^ cofactor was unsupported by the density map in the deposited structure (Fig. 3E). The binding pocket is a general nucleotide binding site^23^, so the AMP molecule favored by EMERALD-ID satisfied the nucleotide restriction while having a better fit into the EM map (Fig. 3F).

### Discovery of unassigned density of deposited EM maps

Given the low resolution of cryoEM maps and lack of ligand identification tools, we suspected that several deposited maps contained regions of density corresponding to ligands that were left unidentified. To remedy this, we searched the EMDB for small molecule-sized unmodeled map regions and screened them using the library of common ligands. Detected regions were filtered by their volume and proximity to the macromolecule so that only the most likely ligand regions were searched. We detected 136 regions from 64 map entries that had a Z-score above -0.5 for the top-ranked ligand, and these entries were further analyzed for identity assignment. Likely identifications are shown in Figure 4.

**Figure 4.**
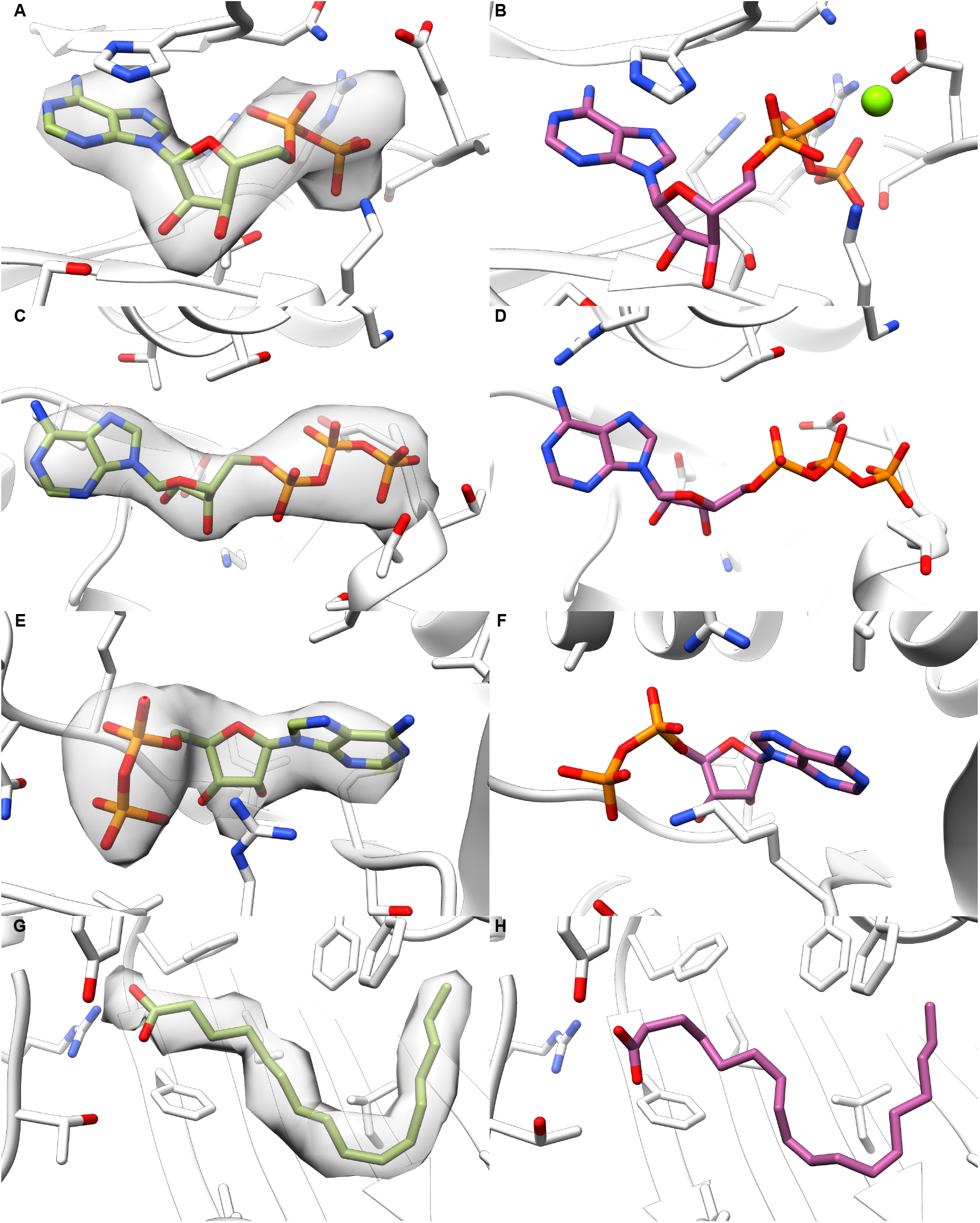
Likely ligands identified in unmodeled regions of deposited maps. (A) Detected ADP molecule in the CLC-7/Ostm1 antiporter (EMDB: 30238, PDB: 7BXU). (B) ATP molecule bound at the same site in (A) in a higher resolution EM structure (PDB: 7JM7). (C) Found ATP molecule in the TRiC complex (EMDB: 33053, PDB: 7X7Y). (D) ATP molecule at an identical site of (C) in a TRiC complex structure from the same study (PDB: 7X3J). (E) ADP molecule ranked first in detected density for a zebrafish Na-K-Cl cotransporter (EMDB: 0473, PDB: 6NPL). (F) ADP bound at nucleotide binding site of a human Na-K-Cl cotransporter (PDB: 7AIQ). (G) Detected palmitate molecule bound to a spike protein of SARS-CoV-2 (EMDB: 11207, PDB: 6ZGI). (H) Linoleic acid bound in the free fatty acid binding pocket in SARS-CoV-2 spike protein (PDB: 6ZB5).

Nucleotide di- and triphosphates were commonly found as unmodeled ligands. In the CLC-7/Ostm1 antiporter^24^, the EM map shows nucleotide-like density, and EMERALD-ID produced an ADP model that fit the map and interacted with the nearby phosphate binding loop (Fig. 4A). This evidence, the confidence of EMERALD-ID, and that ATP was modeled at this site in a higher resolution map^25^ (Fig. 4B) all supported this as a nucleotide binding site. We also identified an ATP molecule at an apparent nucleotide binding site in a known ATPase^26^ (Fig. 4C) that likely went unmodeled because the ligand pocket was not of interest for this protein structure. A structure from the same study modeled ATP at this binding pocket as well (Fig. 4D). Lastly, we found density in a structure of an Na-K-Cl cotransporter in zebrafish^27^ that EMERALD-ID suspected as an ADP molecule (Fig. 4E). Since this structure’s publication, a nucleotide binding site has been determined in the C-terminal domain in the human homolog of the cotransporter^28^ (Fig. 4F), supporting our finding.

Along with nucleotides, our unmodeled density detection found several possible lipid identifications. EMERALD-ID often suggested palmitate molecules in coronavirus spike proteins (Fig. 4G). It is known that spike proteins have fatty acid binding sites^29^ (Fig. 4H), and we previously used EMERALD to model linoleic acid in a spike protein^12^. While it is likely that palmitate is not the exact identity, we detected the signal of a fatty acid binding site nonetheless.

In addition to palmitate molecules, we also found that ten and twelve carbon chain lipids often ranked highly in detected density. For two examples of TRPV channels^30,31^, the density was found in the transmembrane region of nanodisc-reconstituted proteins (Fig. S1) and likely corresponded to disordered lipids from the nanodiscs that cannot be fully identified. While we cannot confidently assign an identity, the frequency of detected regions like these and the abundance of membrane protein structures solved by cryoEM suggest that lipids go undetected in EM maps.

### Identifying uncommon ligands using an endogenous ligand library

While we detected several ligand identities with the common ligand library, we found other density regions that looked like ligands, but evaluation with common ligand identities provided inadequate models. Additionally, microscopists may co-purify an unknown endogenous ligand with a protein sample, which requires a larger ligand library for identification. To cover scenarios that require more ligand identities, we increased the size of the provided library from 30 to 2950 molecules and tested EMERALD-ID’s accuracy on 7 cryoEM structures containing an uncommon ligand.

To determine test cases, we searched the EMDB for entries containing one of the 2950 detected metabolites from the Human Metabolome Database (HMDB)^32^ and looked at rare ligands with 3 or fewer instances in EM structures. After further filtering by ligand size, resolution, and specimen species, we found 14 cases containing an uncommon ligand, which were reduced to 7 after manual inspection. EMERALD-ID ranked the deposited ligand in the top 10% in three out of seven cases (Table 1). For a fourth case (EMDB: 14725, PDB: 7ZH6), the top 10 identities all shared the same steroid core as the endogenous ligand model. Of these four cases, all contained a ligand in the top 5% with a Tanimoto similarity coefficient above 0.75, with three being in the top 1%. Of the cases with low signal for the deposited identity, either the deposited ligand model left unexplained density (EMDB: 34910, PDB: 8HNC) (Fig. S2A) or the ligand signal in the EM map was poor, leading to low Z-scores for all molecules tested (EMDBs: 38692, 38966; PDBs: 8XV5, 8Y65) (Fig. S2B&C).

**Table 1.** Endogenous library screen of uncommon ligand identities.

| Entry | Rank |  | Rank of similar ligand |  |
| --- | --- | --- | --- | --- |
| EMD-30074: HC3 | 7/2701 | (0.3%) | 1/2701 | (0.04%) |
| EMD-25195: BUA | 35/2564 | (1.4%) | 16/2564 | (0.6%) |
| EMD-14725: C0R | 294/2676 | (11.0%) | 15/2676 | (0.6%) |
| EMD-34910: BLR | 629/2753 | (22.8%) | 629/2753 | (22.8%) |
| EMD-35017: GCO | 192/2804 | (6.8%) | 100/2804 | (3.6%) |
| EMD-38692: PXM | 457/2516 | (18.2%) | 457/2516 | (18.2%) |
| EMD-38966: URC | 364/2593 | (14.0%) | 364/2593 | (14.0%) |

Similar ligand defined as an identity with a Tanimoto similarity coefficient greater than 0.75 to the deposited model.

During our search for unmodeled ligands in the previous section, we found instances in the EM map that appeared ligand-like, but none of the common ligands scored well. We decided to screen these regions with the endogenous ligand library to find more probable identity matches. For a Piezo 1 ion channel^33^, a sphingosine lipid was ranked first using the common ligands (Fig. S3A). While the ligand is likely a lipid, the sphingosine model leaves unexplained density, and the density shape and nearby arginine residues suggested a phospholipid identity. Following identification with the HMDB library, the top-ranked molecule was a phosphatidylserine lipid that explains the binding pocket well (Fig. 5A). EMERALD-ID detected a glutamate ligand for an ADH3 structure in *Stenotrophomonas acidaminiphila* ^34^ (Fig. S3B). Another structure of the protein in the same study contains a phenylalanine at this binding site^34^, which was not included in the common ligand library. However, EMERALD-ID detected the amino acid signal, and when a larger library was included, ranked several phenylalanine derivatives within the top 10 structures (Fig. 5B).

**Figure 5.**
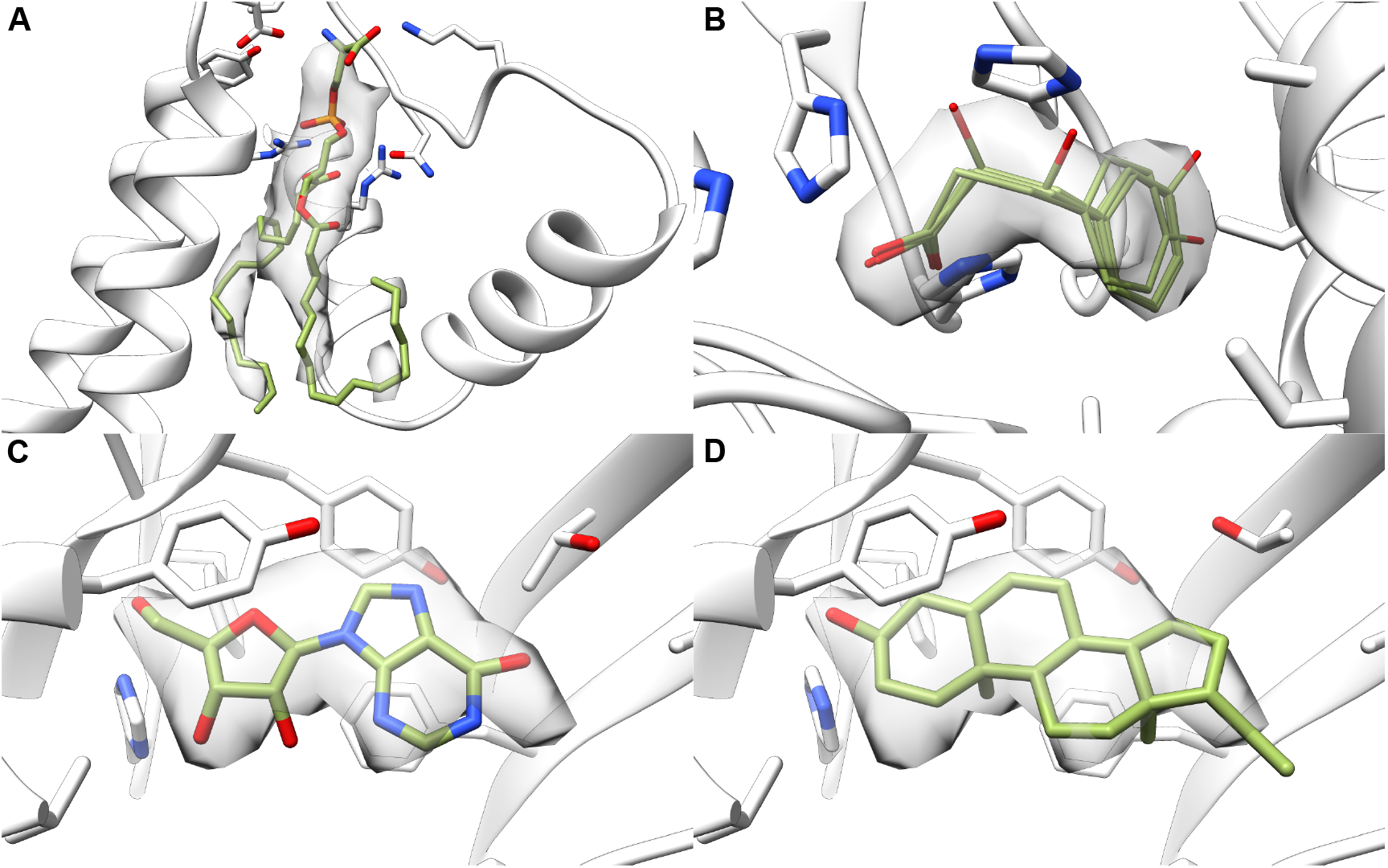
Endogenous ligand search of detected density. (A) Top-ranking phosphatidylserine molecule for detected density in the Piezo 1 ion channel (EMDB: 7128, PDB: 6BPZ). (B) Identities ranked in the top 10 that share features to phenylalanine in ADH3 from *S. acidaminiphila* (EMDB: 35452, PDB: 8IHQ). (C, D) Well-scored identities for detected density at the benzodiazepine binding site in a GABAA receptor (EMDB: 40462, PDB: 8SGO). Inosine (C) ranked fifth overall and allopregnanolone (D) ranked 41^st^.

We also detected a conspicuous ligand blob at the benzodiazepine binding site in a GABA_A_ receptor^35^ (Fig. S3C). Drugs in the benzodiazepines class bind extracellularly to GABA_A_ receptors causing sedative effects, making benzodiazepines effective drugs for anesthetics, seizures, and psychiatric conditions^36–38^. Given the site’s pharmacological importance, endogenous ligands for the site have been sought after, with no known small molecules acting as functional endogenous binders. When we performed an endogenous ligand screen on the detected region, we found 2 plausible identities. Inosine ranked fifth overall (Fig. 5C). Inosine was found to bind to the benzodiazepine site of GABA_A_ receptors^39,40^, but has been discredited as an endogenous binder for weak binding and lack of activity^41^. We also found the neurosteroid allopregnanolone in the top 2% (Fig. 5D). Allopregnanolone was included in the sample preparation of the structure and appeared in the transmembrane region of the deposited model^35^, where it is known to modulate GABA_A_ receptor activity^42,43^. While further experiments will be needed to confirm the ligand identity, EMERALD-ID provided two reasonable explanations of a small molecule bound to the benzodiazepine site.

### Identifying fragments for drug screening experiments

EMERALD-ID proved accurate when distinguishing identities of endogenous ligands, but as cryoEM becomes more relevant for drug discovery^44^, generalizing the method for drug identification becomes crucial. Ligand identification is a necessary task during fragment-based drug discovery. In fragment-based drug discovery (FBDD), low molecular weight molecules that weakly bind to a drug target are determined and used as a scaffold to build a drug candidate. An important step in FBDD is to obtain a structure of a fragment bound to the target. However, the identity of the bound fragment may be unknown if a cocktail of fragments is included during sample preparation. Traditionally, structure determination for FBDD has been achieved through X-ray crystallography because high-resolution is needed to resolve the identity of the ligand. But, many drug targets contain transmembrane regions, precluding the use of X-ray crystallography for their structure determination. As resolution limits improve in cryoEM, it is possible to obtain EM maps with resolvable fragment density — opening FDDD to drug targets that are difficult to solve with X-ray crystallography.

Principles of FBDD were successfully applied to determine high-resolution fragment bound structures by Saur et al.^45^ They resolved 4 structures of fragment-sized ligands bound the cancer target PKM2, two of which included cocktails of 4 fragments during sample preparation. These two structures provided examples for us to test fragment screening with EMERALD-ID. We included both cocktails as libraries for their respective EM maps. EMERALD-ID correctly identified both of the fragments determined by the original authors (Table 2).

**Table 2.** Screening drug fragments for real and simulated EM data.

| Saur et al. |  |  |
| --- | --- | --- |
| Entry (EMDB ID) | Rank | $\Delta G$ Z-score rank |
| EMD-10577: NXE | 1/4 | 3/4 |
| EMD-10584: NXH | 1/4 | 1/4 |
| Simulated data |  |  |
| Entry (PDB ID) | Rank | $\Delta G$ Z-score rank |
| 1EQG: IBP | 1/236 | 1/236 |
| 1GWQ: ZTW | 1/237 | 1/237 |
| 1N1M: A3M | 2/237 | 1/237 |
| 1QWC: 14W | 2/237 | 25/237 |
| 1S39: AQO | 1/232 | 1/232 |
| 1YZ3: SKA | 36/236 | 1/236 |
| 2OHK: 1SQ | 25/236 | 42/236 |

While these results are promising, more examples will be needed to evaluate EMERALD-ID’s utility for FBDD. To provide more test cases at lower resolution, we turned to realistic simulation EM data (details in Methods). We found 7 high-resolution structures solved by X-ray crystallography that contained fragment-sized ligands and simulated EM data for them at 3.5 Å resolution. The native fragments were screened against a combined 238 fragment library from the Cambridge^46^ and York^47^ 3D libraries. For five of the seven entries, EMERALD-ID ranked the native fragment within the top 5 structures (Table 2). Moreover, fragments ranked above the native fragments share similar characteristics to the native fragment (Fig. S4). Despite binding weakly to their receptors, the binding affinity Z-score was powerful at discerning between identities, with 5/7 native fragments ranking first by this metric (Table 2). This suggests that EMERALD-ID can be used for fragment screening, even when the fragment density is poor.

## DISCUSSION

Here, we introduce EMERALD-ID to assign identities to ligand density in cryoEM data. We correctly identified ligands in over 40% of instances that contained a common ligand, a rate much higher than Phenix ligand identification. The power of EMERALD-ID was further shown by identifying several ligands that were left unmodeled during the original deposition. Finally, EMERALD-ID proved effective in plausible scenarios of screening a large endogenous ligand library and a fragment library for drug discovery.

Along with the predicted probability, we believe the values of the Z-scores will be useful when evaluating ligands. We find that 61% of entries in the common ligand benchmarking set have a top scoring identity with a Z-score of -1.0 or greater. Meanwhile, only 6% of our detected unmodeled ligand regions found a ligand identity better than this threshold. This suggests that the Z-score is sensitive to whether a ligand is present in the structure. Additionally, both binding affinity and density Z-scores should be above -1.0 to eliminate ligands that overfit to the map or ignore the map. By calculating these standardized density fit and energy terms, our Z-score metrics could also be valuable in determining the quality of ligand models.

As noted, the success of identification with EMERALD-ID greatly depended on the success of docking the small molecules into density, and the limitations of the method mainly lie with limitations in our ligand docking. Molecules containing inorganic elements like metal ions cannot be properly parameterized for our ligand docking, leaving out ligands like hemes from analysis. Another exclusion from analysis are glycans which, due to their unique structural characteristics and covalent binding, require special methods for docking and identification. Additionally, EMERALD can only dock a single ligand conformation at once, so pockets with multiple ligands or cofactors must be docked successively.

Incorrect identification may still occur even if the true identity is well-fitted. However, most identity confusion in EMERALD-ID occurred between similar identities (Fig. 2D, Table 1), so even if the true identity is not ranked first or included in the library, a ligand in the same class will likely score well. The binding affinity calculations have a slight bias towards large hydrophobic ligands, and molecules with 10 or fewer heavy atoms can benefit from high density correlations from overfitting. We recommend caution if either scenario describes the top identity and suggest using the Z-score guidelines described above to interpret results.

Improvements to EMERALD-ID will likely come from changes in the force field in Rosetta due to the method’s reliance on binding affinity calculations. As mentioned above, these calculations prefer flexible lipids. Corrections to hydrophobic interactions in Rosetta or other advancements in binding affinity calculations via deep learning will alleviate these issues. While our simple linear regression model is effective in estimating binding affinity and ligand map correlation, the model will likely become more accurate with better training data and addition of predictive features — which should occur with standardization of EM tools for ligand validation^3,48,49^. As presented, EMERALD-ID is effective in identity determination for common modeling scenarios, and we hope that its accuracy and ligand Z-score calculations contribute to improved quality of ligand models for better insights into structural biology.

## RESOURCE AVAILABILITY

### Lead contact

Requests for further information and resources should be directed to and will be fulfilled by the lead contact, Frank DiMaio.

### Materials availability

This study did not generate new materials.

### Data and code availability

- Data used for this study are available at https://doi.org/10.5281/zenodo.14056520, organized by their respective figure or table. These data include source data for each plot and underlying data used to generate them, all docked ligand conformations for structures featured in figures, docked ligand conformations of the deposited and top identities for the common ligand screen, and docked structures for every tested identity for the endogenous ligand and drug fragment screens. Projection stacks, metadata files, and reconstructed maps for simulated EM data for Table 2 are available for download at https://files.ipd.uw.edu/pub/EMERALD-ID/Table2.tar.gz, and the same files for Figure S6 are available at https://files.ipd.uw.edu/pub/EMERALD-ID/Table2FigS3.tar.gz.
- Code for EMERALD-ID, the unmodeled density detector, and the cryoEM density simulation are all available in Rosetta for weekly releases after November 12, 2024. Instructions on how to use them and example scripts used for this manuscript are included in the tutorials file at https://doi.org/10.5281/zenodo.14056520.
- Any additional information required to reanalyze the data reported in this paper is available from the lead contact upon request.

## ACKNOWLEDGMENTS

A.M. and F.D. were supported by a grant from Thermo Fisher Scientific. D.P.F. and F.D. were supported by the National Institute of General Medical Sciences (1R01GM123089-01). G.Z. and F.D. were supported by the Defense Threat Reduction Agency (HDTRA1-22-1-0012). We would like to thank the Institute for Protein Design computing team for maintaining the high-performance computing cluster to run our experiments. We’d especially like to thank Bulat Faezov for setting up Phenix for ligand identification. Thank you to Eddie Pryor, Holger Kohr, John Flanagan, and Maurice Peemen for periodic thoughts and discussions on the method.

## AUTHOR CONTRIBUTIONS

**Andrew Muenks:** Conceptualization, Methodology, Software, Validation, Formal Analysis, Investigation, Writing - Original Draft, Writing - Review & Editing, Visualization. **Daniel P. Farrell:** Software, Writing - Review & Editing. **Guangfeng Zhou:** Software, Writing - Review & Editing. **Frank DiMaio:** Conceptualization, Methodology, Software, Writing - Original Draft, Writing - Review & Editing, Funding acquisition.

## SUPPLEMENTAL INFORMATION INDEX

Document S1. Figures S1-S6 and their legends.

## STAR METHODS

### Method details

#### Ligand parameter generation

The outcome of ligand docking and identification greatly depends on the protonation state and partial charges assigned to the ligand. We recommend using our provided ligand parameters or using the MMFF94 force field^50^ to calculate partial charges. For all experiments, hydrogen atoms were added at pH 7.4 to unprotonated SDF files via openbabel (v. 3.1.0)^51^. Partial charges for the protonated files were calculated with the MMFF94 force field in openbabel. The resulting MOL2 files were then converted to Rosetta-specific ligand residue parameters files for docking. The origin of the unprotonated SDF file depended on the experiment. For model training, an SDF file of the first instance of the ligand in its respective structure was downloaded from the PDB. For the common ligand library, SDF files of the ideal geometries for each ligand were used.

The endogenous ligand library and drug fragment libraries started from SMILES strings. The SMILES for the endogenous ligand library were downloaded from the HMDB and converted into 2D coordinates with openbabel. The 2D coordinates were converted to 3D in openbabel with 3 successive rounds of 3D conformer generation on the slowest setting using a final energy minimization with 2000 steps of the steepest descent algorithm. Drug fragments had their SMILES strings protonated with dimorphite (v. 1.2.4)^52^ at pH 7.4. The protonated SMILES strings were converted to 3D coordinates with openbabel on its default speed and then minimized with 2000 steps of the steepest descent algorithm.

#### Local resolution calculation

One feature used in training and evaluation of EMERALD-ID was the local resolution of the binding pocket. To generate these values for all maps across all experiments, we first calculated local resolution maps with MonoRes from the Xmipp software package (v. 3.22.07.0)^53^ and then calculated average local resolution values for all voxels within 5 Å of the ligand’s center of mass.

Local resolution maps were determined by filtering the deposited EM map with a Gaussian kernel with a sigma of 0.02 times the map dimensions. Voxels in the filtered map with a value above 0.05 times the maximum voxel value were saved to a binary mask for the map, which was then used by MonoRes to create the local resolution map. Voxels with local resolution values of zero were excluded from the average calculation. If the average local resolution in the binding pocket was more than 1 Å lower than the global resolution, then the global resolution was used in place of the local resolution.

#### Linear regression model calculation

To train the linear regression model, we took first instances of ligand identities in EMDB entries where the docked ligand conformation was within 1 Å RMSD of the deposited model in our previous EMERALD manuscript^12^. Since we only looked at ligands with 25 or fewer torsion angles when evaluating EMERALD, we supplemented the training data with entries that had ligand identities with over 25 torsion angles and could be docked within 1.5 Å RMSD of the deposited model. The ligands and surrounding flexible residues of all structures were relaxed in the EM map with a Cartesian minimization in Rosetta and their binding affinities and ligand map correlations were calculated. The relationships between these terms and ligand-map features were probed with a linear regression model in R (v. 4.3.1). We found that the number of heavy atoms in the ligand (*a*) predicted binding affinity with Eq. 1

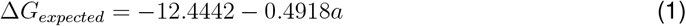

and ligand density correlation could be predicted with the ligand’s heavy atom count, the local resolution of the map 5 Å around the ligand, and the correlation of the entire receptor to the map with Eq.2

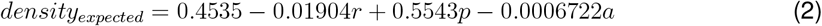

where *r* is the local resolution, *p* is the map correlation of the entire pose, and *a* is the number of heavy atoms,.

The density correlations and binding affinities for all docked identities along with their respective expected values were used to calculate Z-scores where the expected value is the mean and the standard deviation was determined empirically by tuning the standard deviation of the residuals from the linear regression model (*σ*_Δ*G*_ = −10.533, *σ*_*density*_ = 0.043152). Once Z-scores for the binding affinities and density correlations were calculated, they were combined into a single Z-score by averaging the two values and dividing by 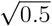. To calculate predicted probabilities, a softmax function was applied to a distribution of modified Z-scores

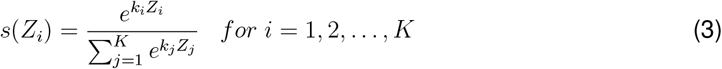

Where **k** is a vector of constants where

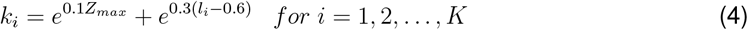

and *Z*_*max*_ is the maximum Z-score of all identities and *l* is the map correlation of the docked ligand identity.

### Determination of common ligand library

We wanted to provide a library of common ligands that can be used for most identification tasks. Common ligand libraries exist in other identification methods^14,15^, but these libraries were created from the entire PDB. Several ligands in the library are ligands relevant for X-ray crystallography but not cryoEM, like cryoprotectants. We decided to create our own library of common ligands specific for cryoEM solved structures. Entries from the EMDB between 2-6 Å global resolution for which a deposited ligand-bound model existed before September 13, 2023 were collected. The first instance of a unique ligand identity in each entry was counted, excluding ligands that cannot be processed by EMERALD, like covalently-bound ligands, ligands containing metal elements, and inorganic compounds. The resulting list of ligands contained several phospholipids. Identities among phospholipids are difficult to parse and require special considerations to dock properly due to their conformational search space, so we excluded examples of phospholipids from the common ligand library. Finally, analogs of higher count common ligands, such as ATP analogs phosphomethyl- and phosphoamino-phosphonic acid adenylate ester, were removed. The remaining ligands with more than 30 instances were separated and provided a library of 30 common ligand identities.

We searched the PDB for EM-solved entries that contain one of the 30 common ligands. Entries were filtered to exclude those with covalently-bound ligands, metal-coordinating bonds, and examples with another small molecule within 10 Å of the ligand of interest. Structures were further excluded if they were missing whole domains modeled into the map. After filtering, we had 1387 entries to screen common ligands with EMERALD-ID.

### Small molecule docking with EMERALD

The EM map, ligand parameter files, and an input model of the receptor were provided to EMERALD for small molecule docking. Input models had all HETATM lines removed except for an ATP model centered on the analyzed ligand blob. The identity provided in the input structure does not matter, as long as the ligand is centered on the density that is being investigated. For each identity in the library, a pool of 100 ligand conformations were generated and optimized over 10 generations of a genetic algorithm as described in the EMERALD manuscript^12^, except when docking drug fragments where a pool size of 50 conformations were used because of their smaller conformational search space. The conformation with the lowest Rosetta energy for each identity was passed to EMERALD-ID for evaluation.

Estimated binding affinity values were calculated using a simple entropy model in Rosetta as described in Zhou et al.^54^ For ligand-map correlation values, we applied a penalty to the value from the EMERALD-docked model because large ligands at low resolutions had ligand map correlations unreasonably high for their fit into the map because of high background density signal from the receptor. The penalty was determined by the difference in map correlation with and without the ligand present (Δlig dens). The penalty was empirically derived by observing cutoffs of Δlig dens values from the training dataset and was calculated with Eq. 5.

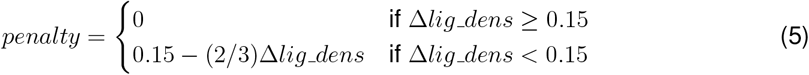

Once calculated, Z-scores were determined as explained above.

EMERALD-ID can be operated sequentially or in parallel, depending on the size of the ligand library. When operating sequentially, the cryoEM map needs to be loaded once and all ligand molecules will be docked in a single job of Rosetta. For large ligand libraries, separate EMERALD runs for each molecule can happen in parallel, and an external python script evaluates and rank all ligands once docking is complete. Examples on how to run in both modes are included in the file repository described in the Code Availability statement.

### Ligand identification of common ligands with Phenix

EM maps for entries in the common ligand dataset were converted to structure factors using phenix.map to structure factors. Ideal CIF files for each ligand in the common ligand library were downloaded from the PDB. The structure factors, an input model without the ligand, a directory containing the CIF files, and a search center of the center of mass of the deposited ligand model were provided to phenix.ligand identification (v. 1.21.1-5286). The rankings of the deposited identity from Phenix identification were compared to EMERALD-ID.

### Unassigned density finder methodology and filtering

The EMDB ligand-bound entries within 2-6 Å global resolution described above along with 3-4 Å maps containing structures without bound ligands were searched for unassigned regions (“blobs”) of the map that could possibly belong to a ligand. We discovered these regions with an unassigned density finding tool. The tool created a mask of the receptor and calculated the mean and standard deviation of all voxel values within the mask. Z-scores for all voxel values outside the mask were calculated using these values. Voxels with a Z-score greater than 0.5 were labeled as peaks and neighboring peak voxels were grouped together to form blobs. Each blob was scored by its number of voxels and fraction of surface voxels that are within 4 Å of the receptor. Blobs were filtered to only keep those with more than 70 voxels, more than 90% of the blob surface interacting with the receptor, and further than 5 Å from a cut or terminus in the protein structure. All blobs passing the filters were screened with the common ligand library with the binding pocket centered on the found blob. After screening, top-ranked ligand identities with a Z-score greater than -0.5 were manually analyzed.

### Endogenous ligand library screening

We obtained 3030 SMILES strings for all detected and quantified metabolites with an endogenous origin from the Human Metabolome Database^32^. After processing as described above, we had 2950 ligand identities to use for docking. Using the RCSB REST API^55^, we searched for rare occurring identities that had one to three cryoEM solved structures containing the respective ligand’s SMILES string. Entries were filtered to those that had a human source organism, a resolution worse than 3.3 Å, and more than 5 heavy atoms in the ligand. This left us with 14 entries, which were then manually pruned to 7 after removing entries with multiple ligands in the binding pocket and large lipids with inconclusive support in the EM map. The 2950 endogenous ligand identities were docked for each of the 7 entries and ranks were determined. Ligand similarity among the endogenous library was calculated by Tanimoto similarity coefficient of small molecule fingerprints with RDkit (release 2024.03.4)^56^.

### Fragment screening preparation

Both examples from the fragment cocktail experiments for pyruvate kinase 2 from Saur et al.^45^ were used for fragment screening. Fragment parameters were created from SMILES strings as described above and a library of the respective cocktails were provided for identification for EMERALD-ID. For examples to use with simulated data, fragment bound crystal structures were taken from Congreve et al.^57^ The Cambridge^46^ and York^47^ 3D libraries provided 137 and 106 fragments, respectively, for the simulated data fragment screening. We chose these libraries because their SMILES strings were publicly available and the fragment sizes in the library were similar to the fragments in the crystal structures.

The sim cryo tool^58^ in Rosetta was used to simulate cryoEM maps for the fragment bound crystal structures. Briefly, sim cryo creates 2D projections of protein structures for map reconstruction rather than attempting to directly simulate the 3D map. The input structure is randomly rotated for a selected number of rotations, and projection images across each XYZ plane are recorded for each rotation. Gaussian noise is applied to each image, and pixels in the projections, which correspond to atoms in the structure, are randomly perturbed to simulate atom heterogeneity.

We simulated cryoEM maps for the crystal structures to a resolution around 3.5 Å by using a Gaussian noise multiplier of 0.6, a pixel size of 1, and an atom perturbation factor of the atom’s B-factor divided by 120 to produce an image stack of 45000 projections. The image stacks of the perturbed projections were passed into cryoSPARC (v.4.4)^59^ and 2D class averages were created. 3D maps were created from 2D classes with ab initio reconstruction and then a homogeneous refinement. Two cases, 1FV9 and 2JJC, could not produce realistic cryoEM maps because of their small size, but all other structures produced simulated maps with realistic low-resolution ligand binding sites (Fig. S5). To further show the quality of simulated data with sim cryo, we simulated EM data of cryoEM-solved structures using the same simulation protocol described above. We found that map correlations for ligand models and their binding pockets were similar for the real EM and simulated EM data (Fig. S6).

### Ligand and data visualization

Figures of ligand-bound models and their EM maps were created using UCSF Chimera (v. 1.17.3)^60^. Plotting of data was performed using the ggplot2 package (v. 3.4.3) in R^61^.

