## Supplemental Information for "Automated identification of small molecules in cryo-electron microscopy data with density- and energy-guided evaluation"

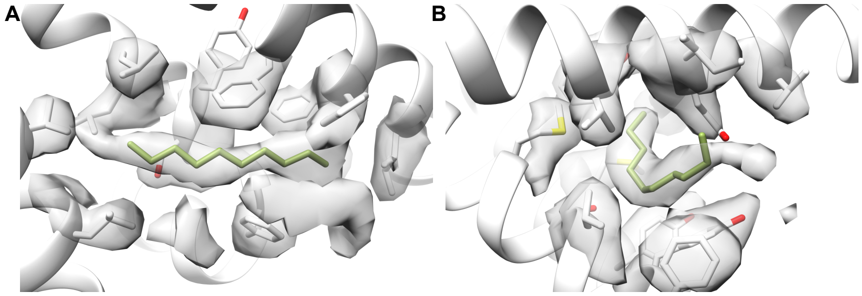


Figure S1. Lipids detected in unmodeled density regions. (A) Ten carbon chain in transmembrane region of TRPV2 (EMDB: 33774, PDB: 7YEP). (B) Ten carbon chain in transmembrane region of TRPV1 (EMDB: 23128, PDB: 7L2H).

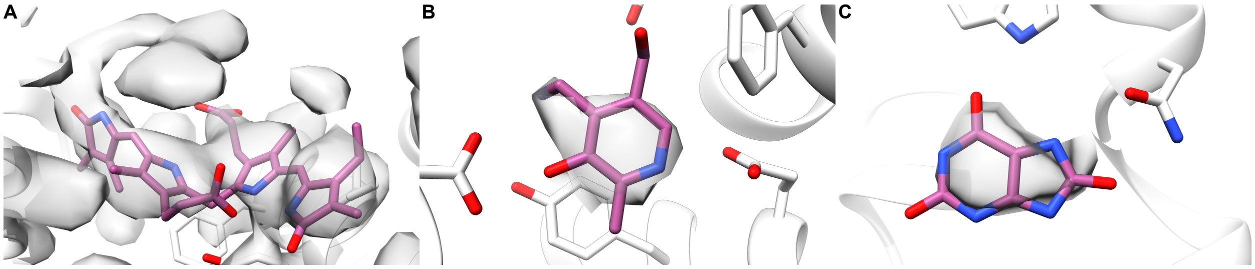


Figure S2. Deposited ligand models for endogenous ligand screens without a similar ligand in the top 10 percent. (A) Bilirubin bound to OATP1B1 (EMDB: 34910, PDB: 8HNC). (B) Pyridoxamine bound to SLC19A3 (EMDB: 38692, PDB: 8XV5). (C) Urate bound to GLUT9 (EMDB: 38966, PDB: 8Y65).

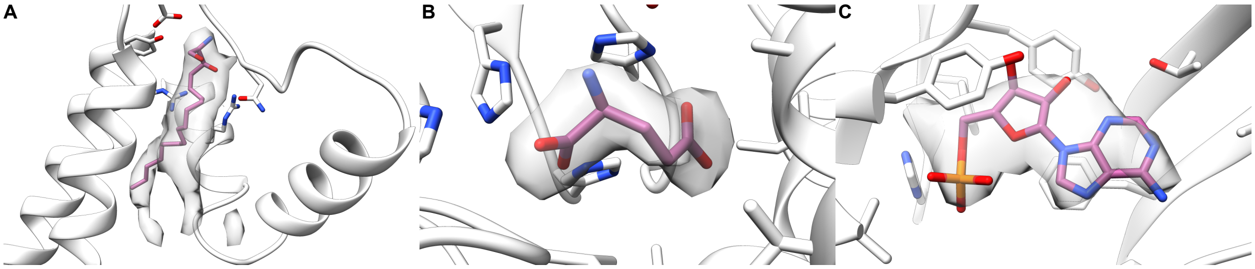


Figure S3. Top common ligand identity for detected density. (A) Detected sphingosine lipid in the Piezo 1 ion channel (EMDB: 7128, PDB: 6BPZ). (B) Detected glutamate molecule in ADH3 from S. acidaminiphila (EMDB: 35452, PDB: 8IHQ). (C) Detected AMP ligand in a $GABA_A$ receptor (EMDB: 40462, PDB: 8SGO).

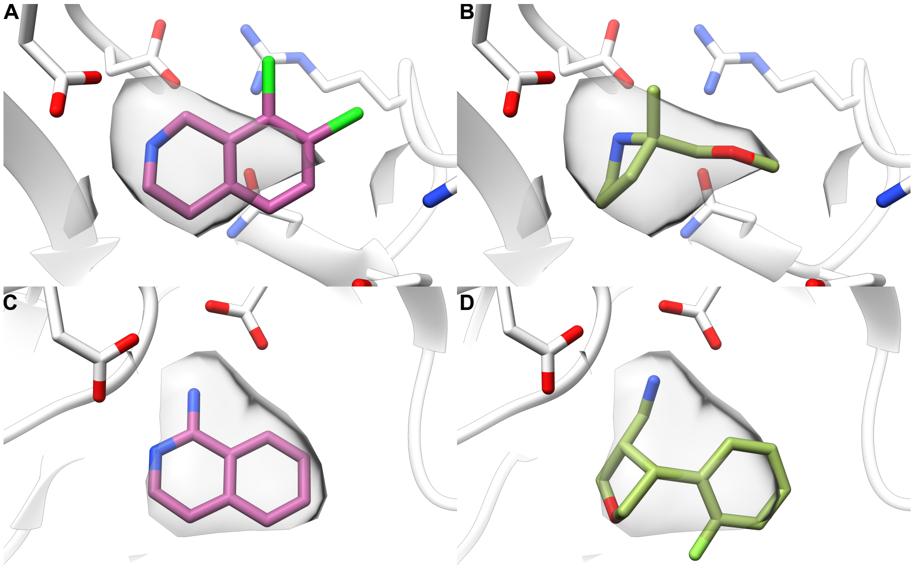


Figure S4. Deposited and top ligand identities for failed fragment screening cases. (A) Deposited and (B) top ranked fragment for a simulated EM map of phenylethanolamine N-methyltransferase (PDB: 1YZ3). (C) Deposited and (D) top ranked fragment for a simulated EM map of beta-secretase (PDB: 2OHK).

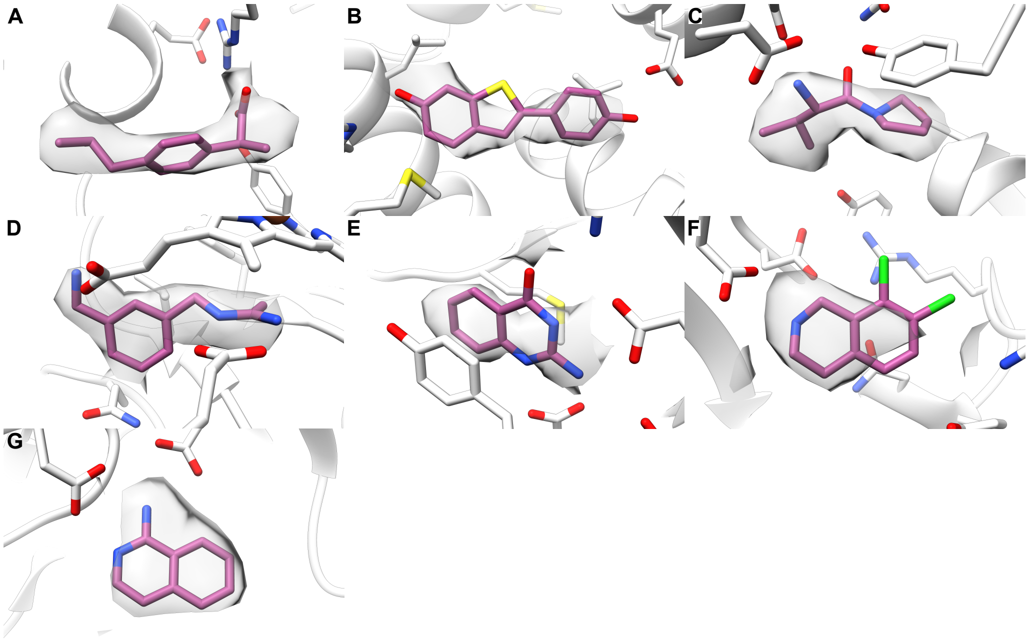


Figure S5 Deposited ligands with their simulated EM maps. (A) Ibuprofen bound to COX-1 (PDB: 1EQG). (B) Raloxifene core bound to estrogen receptor alpha (PDB: 1GWQ). (C) Fragment inhibitor bound to dipeptidyl peptidase IV/CD26 (PDB: 1N1M). (D) W1400 inhibitor bound to nitric oxide synthase (PDB: 1QWC). (E) Fragment inhibitor bound to tRNA-guanine transglycosylase (PDB: 1S39). (F) Fragment inhibitor bound to phenylethanolamine N-methyltransferase (PDB: 1YZ3). (G) 1-amino-isoquinoline bound to beta-secretase (PDB: 2OHK).

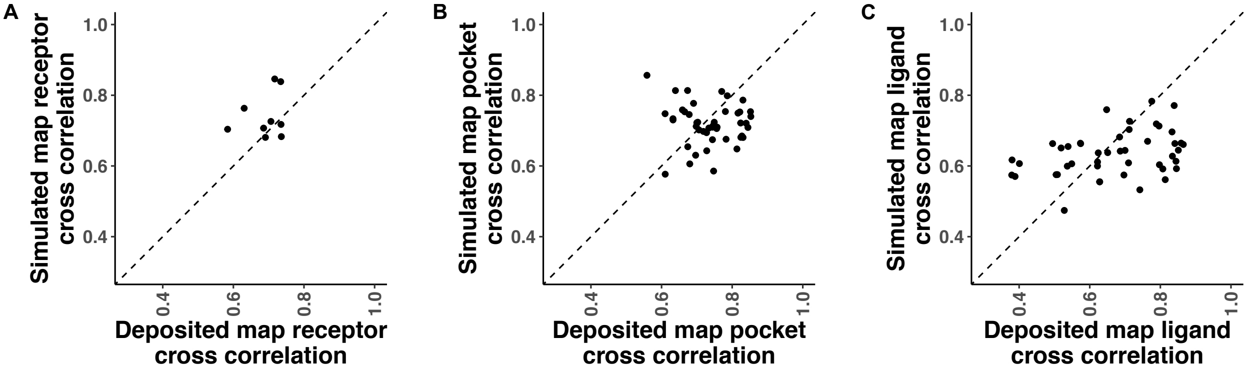


Figure S6. Comparison of select deposited EM maps to their simulated data. (A) Comparison of cross correlation of the receptor to deposited and simulated density maps for 9 EMDB entries. (B) Comparison of cross correlation of the ligand binding pocket to deposited and simulated density maps. Each point corresponds to a small molecule residue of the 9 maps. Pocket is defined as 10 Å around the center of mass of the ligand. (C) Comparison of cross correlation of the ligand model to deposited and simulated density maps for 9 EMDB entries. Each point corresponds to a small molecule residue of the 9 maps.
